# Tazobactam selects for multidrug resistance

**DOI:** 10.1101/2023.08.15.553388

**Authors:** Emma R. Holden, Muhammad Yasir, A. Keith Turner, Ian G. Charles, Mark A. Webber

**Affiliations:** Quadram Institute Bioscience, Norwich Research Park, Norwich, Norfolk, NR4 7UQ, U.K.; Norwich Medical School, University of East Anglia, Norwich Research Park, Norwich, Norfolk, NR4 7TJ, U.K.

**Keywords:** Functional genomics, transposon mutagenesis, TraDIS, efflux, AcrAB, AroK

## Abstract

Piperacillin-Tazobactam is a β-lactam/β-lactamase inhibitor combination which is amongst the most prescribed antimicrobials in hospital medicine. Piperacillin is inactivated by commonly carried resistance enzymes, but tazobactam inhibits these allowing successful treatment. The effect of piperacillin on Gram-negative bacteria has been widely studied, but less attention has been paid to the effects of tazobactam. We used a massive transposon mutagenesis approach (TraDIS-*Xpress*) to determine the genes in *Escherichia coli* that affect survival when exposed to piperacillin and tazobactam, separately and together. We found significant differences in the selective pressure of the two drugs: a striking finding was that multiple efflux pump families and regulators were essential for survival in the presence of tazobactam, but only one efflux system was beneficial for piperacillin. Additionally, we identified the shikimate kinase AroK as a potential target for tazobactam. This method also found that genes involved in DNA replication and repair reduced *E. coli* susceptibility to a combination of piperacillin and tazobactam, not seen from either drug treatment alone. Treatment of *E. coli* and *Klebsiella pneumoniae* with piperacillin and/or tazobactam selected for mutants with reduced susceptibility, and SNP analyses supported the TraDIS-*Xpress* findings that tazobactam selects for changes in membrane permeability and maintenance associated with multidrug-resistance. Increased efflux activity is an important foundation of multidrug resistance in human pathogens, therefore the finding that tazobactam can select for this is concerning. These findings could have consequences for antibiotic prescription and should inform the development of future β-lactamase inhibitors to reduce the global increase in multidrug-resistant infections.

## Introduction

Piperacillin-Tazobactam is one of the most commonly prescribed antibiotic treatments in the UK, used to treat a wide range of infections including pneumonia, urinary tract infections and skin and soft tissue infections (Perry and Markham, 1999). Beta-lactam antibiotics such as piperacillin disrupt peptidoglycan synthesis, an essential component of the bacterial cell envelope. Resistance to this class of antibiotic can however be conferred by genes coding for β-lactamases which cleave the β-lactam ring, rendering the antibiotic ineffective. A wide variety of extended-spectrum β-lactamases have been described and have increased in prevalence since the first was described in 1983 (Knothe et al., 1983). These can be chromosomal but are often spread on conjugative plasmids and are now common in many species including those causing both hospital-acquired and community-acquired infections (Cantón et al., 2008). To prevent the activity of β-lactamases, β-lactam antibiotics can be given in conjunction with a β-lactamase inhibitor, such as Tazobactam. These inhibit β-lactamase enzymes so that activity of the antibiotic is protected. Extensive research has been conducted on how the cell responds to treatment with β-lactam antibiotics, including piperacillin (Kong et al., 2010). However, relatively little work has focussed on how bacteria respond to β-lactamase inhibitors, including tazobactam and whether these compounds have any significant activity beyond their interactions with β-lactamases (Tooke et al., 2019).

Large-scale genomic screens have previously been used to identify the extended complement of genes and pathways that affect susceptibility to a given antibiotic, including genes beyond the principal target. These methods have often revealed many genes contribute to sensitivity to antibiotics to varying degrees. We have used one such method, TraDIS-*Xpress*, where massively dense transposon mutant libraries investigate bacterial loci involved in responses to stress at almost base pair resolution. These libraries make use of a transposon-encoded outward-transcribing inducible promoter, which allows changes in gene expression as well as gene disruption to be assayed for roles in survival under a given stress. This provides information about essential genes, which are often those of interest for antibiotics that usually target essential genes. This approach has recently been used to identify known and unknown mechanisms of action and resistance to triclosan (Yasir et al., 2020), fosfomycin (Turner et al., 2020b), fluoroquinolones (Turner et al., 2020a), meropenem (Thomson et al., 2022), trimethoprim and sulfamethoxazole (Turner et al., 2021).

In this study we used TraDIS-*Xpress* to identify how *Escherichia coli* responds to piperacillin and tazobactam, separately and in combination. We identified known genes involved in sensitivity to piperacillin but also found many genes under selective pressure after exposure to tazobactam. These included a suite of genes involved in multiple drug resistance with multidrug efflux pumps and regulators, whereas only one efflux system was implicated in piperacillin susceptibility. To explore the implications of these findings, we exposed *E. coli* and *Klebsiella pneumoniae* to both piperacillin and tazobactam and selected resistant mutants to both. Analysis of these resistant mutants revealed tazobactam selected for mutations in genes involved in efflux activity and regulation and membrane permeability more readily than piperacillin. The demonstration that tazobactam promotes selection of multidrug resistance shows the impact of β-lactamase inhibitors on target bacteria warrants further study.

## Results

### Multiple efflux systems affect susceptibility to tazobactam, but only AcrAB affects piperacillin susceptibility

The TraDIS-*Xpress* data identified 41 genes that affected the susceptibility of *E. coli* to piperacillin, 74 to tazobactam and 108 to a combination of the two drugs, with 159 genes identified in total from all three conditions (supplementary table 1). The variation of sequence reads per insertion site between replicates was low for all conditions tested (supplementary figure 1), indicating a high degree of experimental correlation.

After exposure to tazobactam alone, TraDIS-*Xpress* found many efflux systems affected susceptibility. Increased expression of *acrA, acrB, acrE, acrF, mdtE, mdtF* and *mdfA* appeared to reduce susceptibility to tazobactam, whilst transposon insertions into these genes increased susceptibility (Figure 1). The same could be seen for positive regulators of these systems (*marA, soxS* and *rob*), where increased expression was beneficial to survival and inactivation was detrimental to survival in the presence of tazobactam. Transposon insertions in negative regulators of efflux strongly reduced susceptibility to tazobactam: the log_2_-fold differences in insertions in *acrR* (11.8), *marR* (14.9) and *soxR* (15.3) indicate a very strong selective pressure favouring inactivation of these genes in the presence of tazobactam. In contrast to the tazobactam data, when the library was exposed to piperacillin alone only *acrA* and *acrR* were found to affect fitness with smaller log fold changes in insertion frequency (Supplementary table 1). This data reveals that efflux activity and regulation is extremely important for survival in the presence of tazobactam, but not for piperacillin. When piperacillin and tazobactam were combined, increased expression of *acrA, acrB, acrE, acrF, marA, soxS* and *rob* were beneficial to survival which reflects the combined selective impact of both drugs.

**Figure 1:**
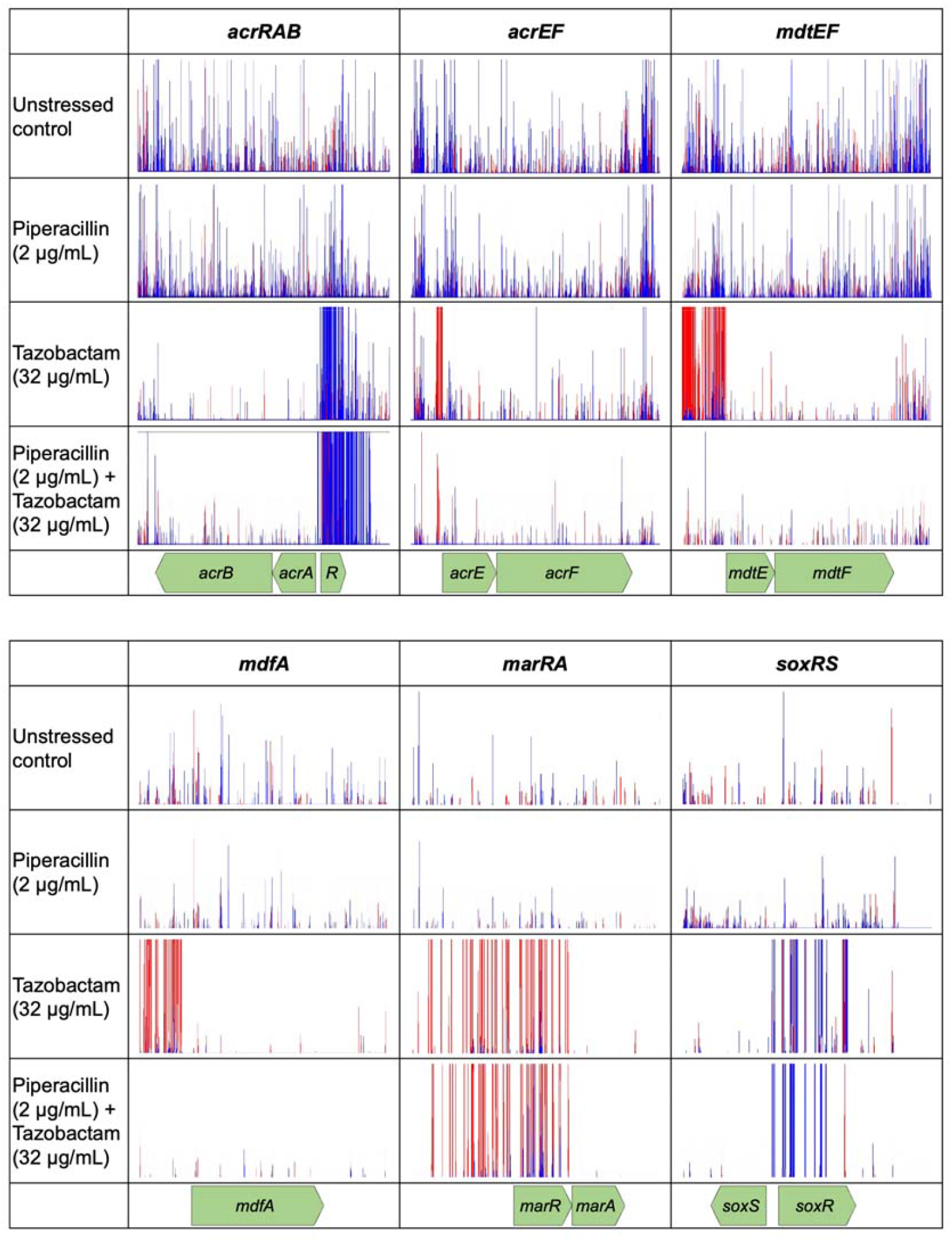
Insertion frequency in and around genes expressing efflux pumps and their regulators in *E. coli* treated with piperacillin and tazobactam, both separately and in combination, relative to an unstressed control. Each fine vertical line indicates the location of a transposon insertion, red indicates the transposon orientation is such that the outward-transcribing promoter transcribes left-to-right and blue right-to-left. Experiments were performed in duplicate, but data from one only is presented due to the high consistency.

Genes with roles in efflux regulation also affected tazobactam susceptibility. The Lon protease is one of these, which reduces the availability of the efflux activators MarA and SoxS. Following growth in tazobactam there was a large (7.4 log_2_-fold) increase in insertion mutations in *lon* compared to controls without tazobactam, indicating that inactivation of Lon increases susceptibility to tazobactam, possibly due to the absence of its protease activity upon MarA and SoxS efflux activators. In the presence of piperacillin alone the opposite was true, and Lon activity reduced susceptibility. This was also seen for genes involved in the synthesis of osmoregulated periplasmic glucans, *opgG* and *opgH*, which increased tazobactam susceptibility and reduced piperacillin susceptibility. Osmoregulated periplasmic glucans have been previously linked to efflux activity (Holden et al., 2023) and are thought to affect the activity of two component signalling systems (Bontemps-Gallo et al., 2017, Bontemps-Gallo et al., 2013), which were also found to have a strong effect on tazobactam susceptibility. The PhoPQ signal transduction system reduced tazobactam susceptibility and the CpxAR system increased susceptibility. These two signalling systems have also previously been implicated in efflux activity and may affect tazobactam susceptibility through this route (Holden et al., 2023). Altogether, we show how the genes involved in efflux regulation and membrane integrity affect survival differently in the presence of each drug, highlighting of the mechanisms through which the cell may tailor its drug response to each drug.

To validate the results of these TraDIS-*Xpress* experiments, *E. coli* was cultured under increasing concentrations of piperacillin, tazobactam and a combination of the two to select for resistant mutants. In the population of mutants selected in the presence of tazobactam, whole genome nucleotide sequencing identified four separate single nucleotide mutations related to efflux (supplementary table 2). These included a frameshift mutation in *acrR* and three distinct mutations in *marR*, both repressors of efflux. In the population exposed to piperacillin alone, one frameshift in *marR* was identified as well as a missense mutation in *rpoD*, which encodes a sigma factor involved in exponential growth (Shimada et al., 2014). To determine if similar selective pressures may be exerted by each drug in other pathogenic bacteria, *Klebsiella pneumoniae* was treated similarly with piperacillin and/or tazobactam to select for resistant mutants. In mutants selected with tazobactam with or without piperacillin, mutations in *acrR* were found again, showing that tazobactam selects for increased efflux pump expression in *K. pneumoniae* (supplementary table 2). Other genes were also mutated including those affecting membrane permeability and integrity. These included the outer membrane porin *ompC* (Cai and Inouye, 2002), *cpxA* involved in regulating membrane integrity (Batchelor et al., 2005), *rfbB* and *rfbD* involved in O-antigen biosynthesis and export (Stevenson et al., 1994) and *ftsI* encoding penicillin binding protein 3 (Suzuki et al., 1978). Together with our findings in *E. coli*, this demonstrates that tazobactam can select for changes in efflux and membrane permeability in two distinct human pathogens.

### The shikimate kinase AroK had a strong effect on growth and survival in the presence of tazobactam in *E*. *coli*

Analysis of the TraDIS-*Xpress* data found a 10.6 log_2_-fold increase in insertion mutations in *aroK* in conditions treated with tazobactam relative to control conditions without tazobactam (figure 2), indicating that inactivation of *aroK* was strongly beneficial for survival in the presence of tazobactam. Supporting this finding, we also found a frameshift variant of *aroK* in an *E. coli* mutant continuously exposed to increasing concentrations of piperacillin and tazobactam (Table 1). AroK is a shikimate kinase involved in chorismate biosynthesis, an intermediate in the synthesis of phenylalanine, tyrosine and tryptophan and also a precursor of folic acid, ubiquinone, menaquinone, and enterochelin (Vinella et al., 1996). AroK is one of two shikimate kinases in *E. coli* alongside AroL. We did not see a significant difference in transposon insertion frequency within *aroL* in the TraDIS-*Xpress* data after tazobactam exposure compared to controls. Deletion of *aroK* or *aroL* did not significantly affect tazobactam susceptibility in *E. coli* (data not shown).

**Figure 2:**
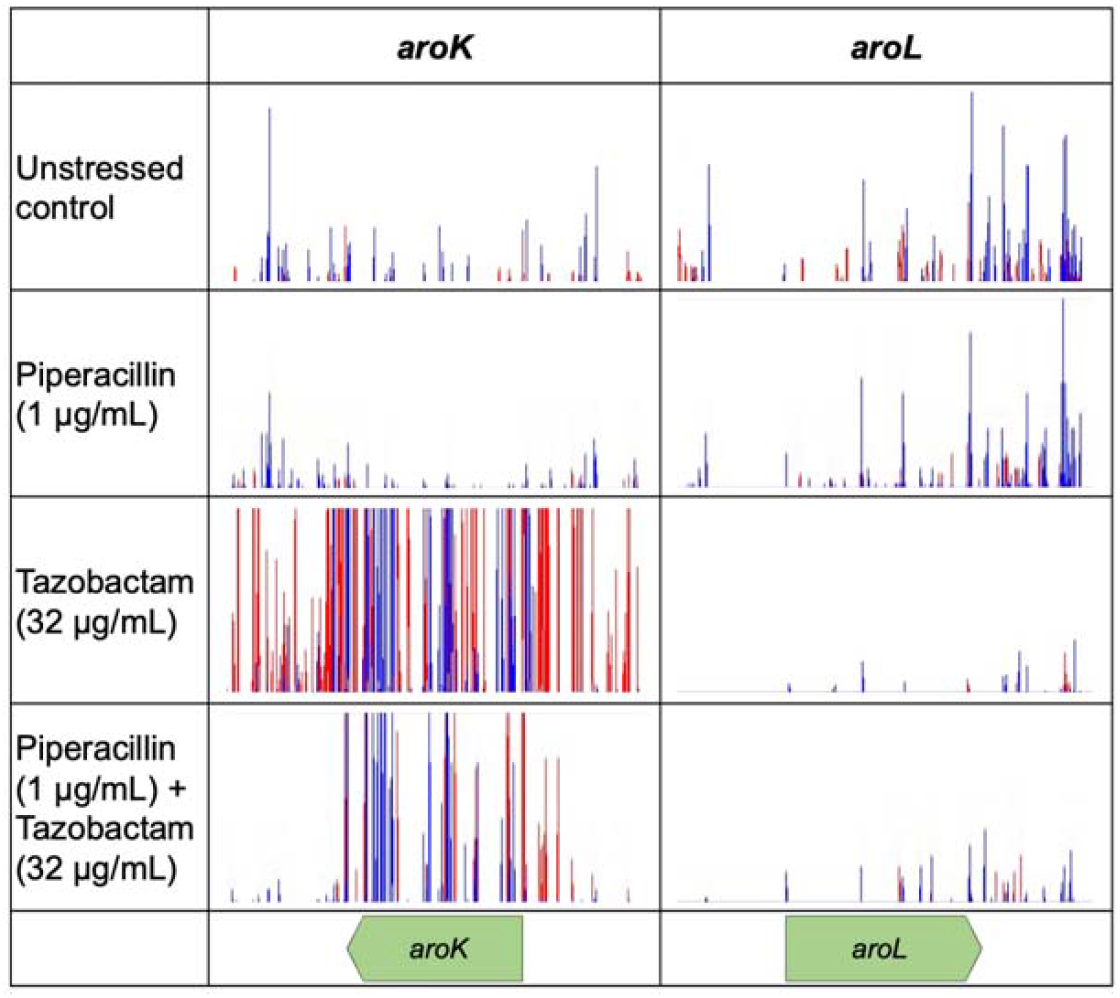
Insertion frequency in and around *aroK* and *aroL* in *E. coli* treated with piperacillin and tazobactam, both separately and in combination, relative to unstressed controls. Red lines indicate the transposon-located promoter is facing left-to-right and blue lines show the promoter facing right-to-left. Images are representative of two independent replicates.

### Genes involved in DNA replication, transcription and translation have a common effect on survival independent of drug treatment

There were very few similarities in the genes that affected susceptibility of *E. coli* to piperacillin and tazobactam: the majority of these similarities came from those involved in translation, where their inactivation reduced susceptibility in all conditions relative to unstressed controls. These included genes involved in tRNA modification and fidelity (*truA* (Kammen et al., 1988), *mnmE* and *mnmG* (Brégeon et al., 2001)), ribosome biogenesis, (*bipA* (Choudhury and Flower, 2015)), as well as *dsbA*, responsible for forming disulphide bonds in periplasmic proteins (Bardwell, 1994). We also found *rapA*, important for recycling RNA polymerase (Sukhodolets et al., 2001), reduced susceptibility to piperacillin and tazobactam, both separately and in combination when inactivated. These findings suggest a reduction in transcription rate and translation fidelity may benefit survival to a wide range of stresses.

We identified that DNA repair and replication reduces susceptibility to both piperacillin and tazobactam, but not each drug separately. Increased expression of *uspB* involved in DNA recombination repair, and *mutT* involved in maintaining DNA replication fidelity, were both beneficial for survival in the presence of both drugs and not each drug separately (Figure 3). *XseB*, encoding an exonuclease subunit involved in DNA repair, was essential to survival in the presence of both drugs, where transposon mutants with an inactive copy of *xseB* could not be detected. This suggests the combination of piperacillin and tazobactam elicits a DNA damage response from the cell and genes involved in DNA repair confer increased fitness when exposed to both drugs, but not either drug separately. This highlights the synergy between piperacillin and tazobactam independent of β-lactamase inhibition.

**Figure 3:**
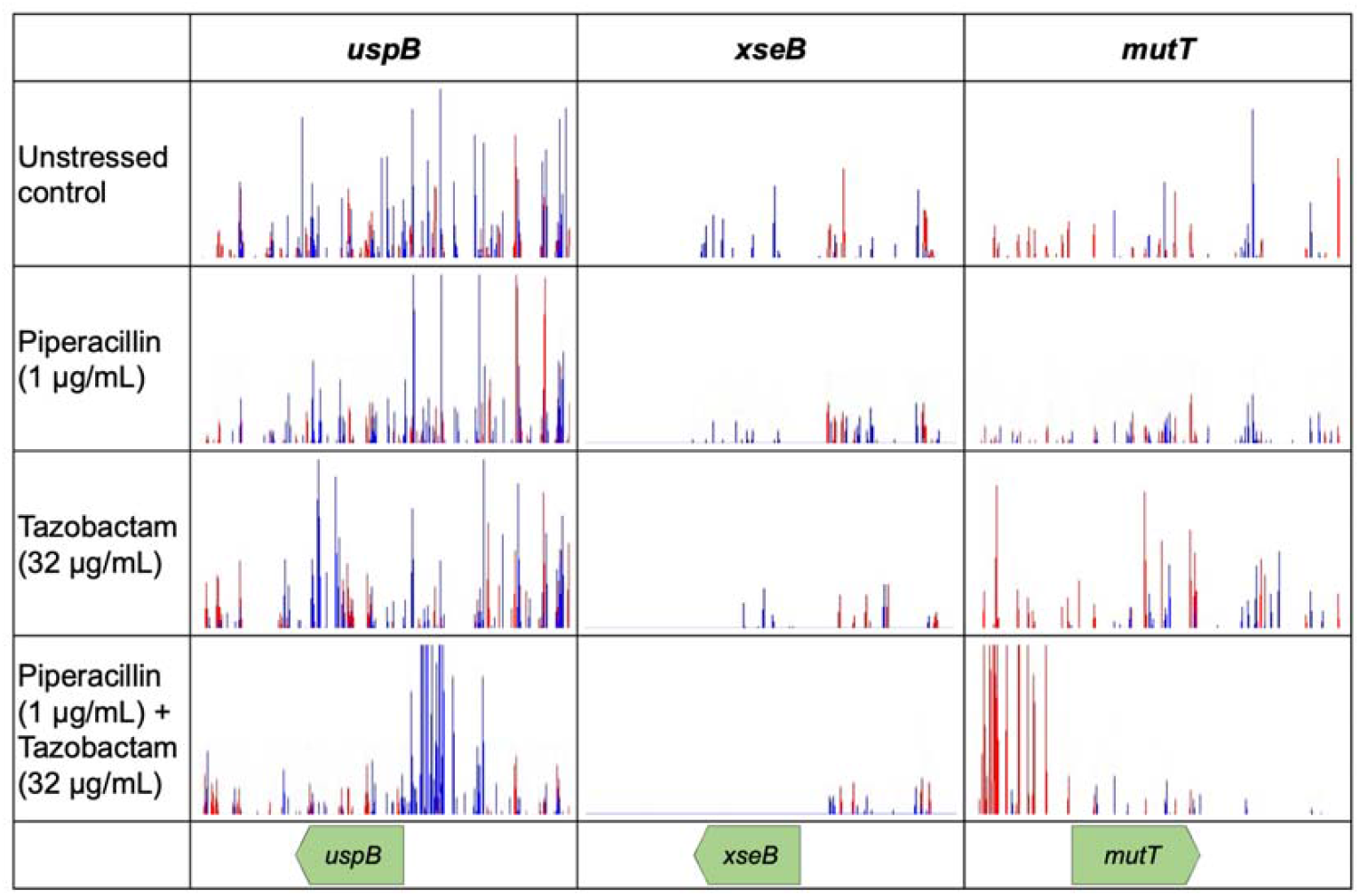
Insertions around genes involved in DNA repair and replication in *E. coli* treated with piperacillin and tazobactam, both separately and in combination, relative to unstressed controls. Red lines indicate the transposon-located promoter is facing left-to-right and blue lines show the promoter facing right-to-left. Images are representative of two independent replicates.

### A range of other genes in diverse pathways were also under selective pressure after exposure to piperacillin or tazobactam

Identification of genes known to affect piperacillin susceptibility confirms the efficacy of this approach. Both *mrcB* and *lpoB* (both involved in peptidoglycan synthesis) were beneficial for survival in piperacillin-exposed conditions. Mutations in genes involved in peptidoglycan synthesis and re-modelling, including *ldcA, mepS, nlpI, prc, slt* and the Tol-Pal system, also demonstrated increased susceptibility to piperacillin both with and without tazobactam, indicating that their encoded functions are beneficial in the presence of piperacillin. The genes involved in the synthesis and function of penicillin binding protein 1A, *mrcA* and *lpoA*, were only beneficial to survival in conditions treated with the combination of both drugs, not each drug separately. This demonstrates the different selective pressure on genes coding for cell wall functions when these drugs are given in combination relative to separately.

We found mutations inactivating glutathione synthesis via *gshA* and *gshB* strongly reduced tazobactam susceptibility. Glutathione has previously been linked to efflux activity (Song et al., 2021, Holden et al., 2023), so this further supports our hypothesis that tazobactam elicits a strong efflux response in *E. coli*.

Multiple genes involved in Fe-S cluster synthesis and repair affected tazobactam susceptibility in *E. coli*. TraDIS-Xpress found inactivation of *hscA* and *hscB* (Tokumoto and Takahashi, 2001) reduced susceptibility and inactivation of *iscS* and its regulator *iscR* (Schwartz et al., 2001) increased susceptibility. IscS also has a role in tRNA modification, however the genes downstream in this pathway (*tusA, tusBCD, mnmE, mnmG*) (Ikeuchi et al., 2006) were found to have a negative effect on survival in tazobactam-exposed conditions. Production of extracellular polysaccharides was also beneficial for survival in piperacillin-exposed conditions. Genes involved in LPS biosynthesis (*wbbK, waaS, lpxL*) and transport (*lptA, lptB* and *lptC*) and most of the operon involved in enterobacterial common antigen (ECA) biosynthesis (*wecA-wecG*) were found to positively affect survival in the presence of piperacillin.

## Discussion

By screening every gene in the *E. coli* genome, we aimed to dissect the differences in genes under selective pressure after exposure to piperacillin, tazobactam, or a combination of the two. We found that tazobactam elicited a strong efflux response from a wide variety of efflux pump families, their local and global regulators, as well as genes that affect global regulator activity. This was in stark contrast to the much more limited efflux response elicited by piperacillin, which consisted of only the AcrAB efflux pump and its local regulator AcrR. When cells were treated with a combination of piperacillin and tazobactam, the global regulators MarA, SoxS and Rob and an additional efflux pump AcrEF were beneficial for survival showing the selective impact of tazobactam is maintained when combined with piperacillin. We also showed that tazobactam readily selects for efflux mutants in *E. coli* and *K. pneumoniae*, two pathogens where efflux mediated resistance is important. Mutations analogous to those we selected within efflux regulators *marR* and *soxR* are well characterised and have previously been described to reduce susceptibility to multiple antibiotics in pathogens (Piddock, 2006). Whilst efflux activity alone may only confer modest reductions in susceptibility to antibiotics it synergises with other resistance mechanisms. For example, *E. coli* that are resistant to fluoroquinolones through mutations in *gyrA* are rendered susceptible by disrupting efflux activity (Kern et al., 2000, Oethinger et al., 2000). This demonstrates that a β-lactamase inhibitor which is used to nullify a resistance mechanism may in fact be actively promoting the emergence of another multidrug resistance mechanism. Usage of Piperacillin and Tazobactam is increasing rapidly (Cotteret et al., 2016, Drawz and Bonomo, 2010), as is global antimicrobial resistance (O’Neill, 2014), so it is extremely important to understand how the antibiotics we use may contribute to resistance.

Tazobactam has been well characterised as an inhibitor of β-lactamases but little previous work has studied any other impacts on the bacterial cell. In addition to the evidence for tazobactam to be selective for increased efflux, we also found a very strong signal for interaction with the shikimate kinase AroK. This may suggest a direct interaction with tazobactam and AroK as insertional inactivation of *aroK* considerably increased growth and survival of *E. coli*, and a SNP within *aroK* was found in an *E. coli* mutant selected after exposure to piperacillin-tazobactam. Shikimate is a precursor for the synthesis of aromatic amino acids, enterobactin, folate, ubiquinone and menaquinone: it is possible that reduced synthesis of these compounds slows respiration, carbon metabolism and therefore growth in order to become tolerant to tazobactam. However, we only see a signal for *aroK* and not *aroL* with the same function, so it is unlikely that this is the mechanism through which reduced tazobactam susceptibility is achieved. AroK has been previously implicated in mecillinam resistance, where deletion reduced susceptibility independently of its shikimate kinase activity suggesting interactions with β-lactams are possible (Vinella et al., 1996). It is possible that AroK affects tazobactam susceptibility in the same manner. It is also possible that the combination of reduced AroK activity and increased efflux activity is responsible for a significant reduction in susceptibility to tazobactam. Further investigation into how tazobactam selects for reduced susceptibility in strains without functional β-lactamases is crucial. If tazobactam can select for loss of function of *aroK*, this could have implications for its susceptibility to mecillinam and other β-lactam antibiotics.

As well as the impacts on efflux and *aroK*, we also found many other genes under selective pressure from either tazobactam, piperacillin or the combination of the two. Previous work by Valéria dos Santos et al. (2010) characterised an *E. coli* isolate resistant to piperacillin and tazobactam and found changes to a diverse group of proteins including upregulation of those in membrane permeability and DNA stress response and a lower abundance of proteins involved in respiration and translation. Our work also found genes involved in DNA replication and repair were beneficial to survival in conditions exposed to both piperacillin and tazobactam, but not each drug separately. This and previous work suggest the combination of both drugs induces DNA damage. We also identified multiple genes involved in replication, transcription and translation affected fitness when exposed to piperacillin and tazobactam. Mutations in genes coding for these functions may slow or stop growth, resulting in increased tolerance to the antibiotics in a non-specific manner as has been shown in previous studies (Brauner et al., 2016)

The demonstration here that tazobactam exerts a selective pressure has implications for how future β-lactamase inhibitors are designed. It is extremely important to prevent selection for multidrug resistance which could be an unintended correlate impact from tazobactam use. Multiple novel β-lactam/β-lactamase inhibitor combinations are in development (Naas et al., 2017, Boyd et al., 2020) and our work shows consideration of the selective impact of the inhibitor should not be overlooked. It remains to be seen if other β-lactamase inhibitors will have significant selective impacts of their own and this should be an important consideration for development of new combinations which are urgently needed to counter the emergence of resistance.

## Methods

### Bacterial strains and growth conditions

The *E. coli* BW25113 transposon mutant library used in this study contains a pool of over 800,000 different mutants and has been described by (Yasir et al. (2020). Approximately 10^7^ CFU/mL of this library was added to 1 mL LB Miller broth in a 96-well deep-well plate and supplemented with concentrations of piperacillin around the MIC (MIC was 4 μg/mL piperacillin, concentrations used were ¼x, ½x, 1x and 2x MIC). Tazobactam was either added or omitted from these conditions at a concentration of 32 μg/mL. The Tn*5* transposon used to create this library contains an outward-transcribing *tac* promoter, which was either induced with 0.2 or 1 μM IPTG or left uninduced. Each condition was carried out in duplicate with two antibiotic-free controls. Cultures were grown for 24 hours at 37 □C shaking and were centrifuged at 3000 x g for 10 minutes to pellet the cells.

### TraDIS-*Xpress* nucleotide sequencing & Informatics

Genomic DNA was extracted from cell pellets following the protocol described by Trampari et al. (2021) and quantified using a Qubit HS assay kit (Invitrogen). Genomic DNA was fragmented with the MuSeek DNA fragment preparation kit (ThermoFisher) and purified using AMPure XP beads (Beckman Coulter). DNA fragments were amplified by PCR using customised biotinylated oligonucleotides of nucleotide sequence specific for hybridisation to one transposon end. Biotinylated DNA fragments were then purified using activated beads from the Dynabeads® kilobaseBINDER™ kit (Invitrogen). Following this, the Dynabeads® with bound DNA fragments were used as the template for a second PCR amplification step, using customised oligonucleotides specific for hybridisation to one transposon end and oligonucleotides specific for the MuSeek adapter nucleotide sequences. Dynabeads® were removed from the PCR reactions using a magnetic stand. The resulting PCR products were purified and fragments of between 300-500 bp in length were applied to a NextSeq 500 using a NextSeq 500/550 High Output Kit v2.5 (75 Cycles) (Illumina).

Output FastQ files from the NextSeq 500/550 were aligned to the *E. coli* BW25113 (CP009273) reference genome using BioTraDIS (version 1.4.3) incorporating BWA (Barquist et al., 2016). Data from conditions treated with different concentrations of IPTG were amalgamated to simplify the data interpretation. The tradis_comparison.R command (part of the BioTraDIS toolkit) was used to determine significant differences (*p* < 0.05, after correction for false discovery) in insertion frequencies per gene between control and test conditions. Increased insertion mutations 5 ‘ to genes, that likely signify genes where increased transcription conferred a selective advantage, were identified visually using the Artemis genome browser (Carver et al., 2011).

### Selection of drug-resistant mutants & SNP analysis

Approximately 10^7^ CFU/mL of *E. coli* BW25113 or *K. pneumoniae* DSM30104 (ATCC 13883) was added to 5 mL LB broth with 1 μg/mL piperacillin and 2 μg/mL tazobactam, either separately or in combination and incubated at 37 □C with shaking at 250 rpm. After 24 hours, 0.1 mL from each condition was transferred into 5 mL fresh LB broth supplemented with piperacillin and tazobactam at double the concentration of the previous condition. At each passage, 1 mL of culture was also collected, pelleted, and stored at -70 □C. Cultures were passaged every 24 hours until no growth was detected in each condition and three independent replicates were performed for each condition. The final passage with visible cell growth was diluted and spread on LB agar to grow single colonies. Three single colonies from each condition were chosen to represent each final population. DNA was extracted from cultures grown from these single colonies and whole genome nucleotide sequence was determined following protocols described by Trampari et al. (2021). FASTQ files generated were compared to the *E. coli* (CP009273) or *K. pneumoniae* (AJJI00000000.1) reference genomes using Snippy version 4.6 (https://github.com/tseemann/snippy) to find SNPs.

## References

Bardwell, J. C. 1994. Building bridges: disulphide bond formation in the cell. Mol Microbiol, 14, 199–205.

Barquist, L., Mayho, M., Cummins, C., Cain, A. K., Boinett, C. J., Page, A. J., Langridge, G. C., Quail, M. A., Keane, J. A. & Parkhill, J. 2016. The TraDIS toolkit: sequencing and analysis for dense transposon mutant libraries. Bioinformatics, 32, 1109–11.

Batchelor, E., Walthers, D., Kenney, L. J. & Goulian, M. 2005. The Escherichia coli CpxA-CpxR Envelope Stress Response System Regulates Expression of the Porins OmpF and OmpC. Journal of Bacteriology, 187, 5723–5731.

Bontemps-Gallo, S., Bohin, J.-P., Lacroix, J.-M. & Slauch, J. M. 2017. Osmoregulated Periplasmic Glucans. EcoSal Plus, 7.

Bontemps-Gallo, S., Madec, E., Dondeyne, J., Delrue, B., Robbe-Masselot, C., Vidal, O., Prouvost, A. F., Boussemart, G., Bohin, J. P. & Lacroix, J. M. 2013. Concentration of osmoregulated periplasmic glucans (OPGs) modulates the activation level of the RcsCD RcsB phosphorelay in the phytopathogen bacteria Dickeya dadantii. Environ Microbiol, 15, 881–94.

Boyd, S. E., Livermore, D. M., Hooper, D. C. & Hope, W. W. 2020. Metallo-β-Lactamases: Structure, Function, Epidemiology, Treatment Options, and the Development Pipeline. Antimicrobial Agents and Chemotherapy, 64, e00397–20.

Brauner, A., Fridman, O., Gefen, O. & Balaban, N. Q. 2016. Distinguishing between resistance, tolerance and persistence to antibiotic treatment. Nature Reviews Microbiology, 14, 320–330.

Brégeon, D., Colot, V., Radman, M. & Taddei, F. 2001. Translational misreading: a tRNA modification counteracts a +2 ribosomal frameshift. Genes Dev, 15, 2295–306.

Cai, S. J. & Inouye, M. 2002. EnvZ-OmpR interaction and osmoregulation in Escherichia coli. J Biol Chem, 277, 24155–61.

Cantón, R., Novais, A., Valverde, A., Machado, E., Peixe, L., Baquero, F. & Coque, T. M. 2008. Prevalence and spread of extended-spectrum β-lactamase-producing Enterobacteriaceae in Europe. Clinical Microbiology and Infection, 14, 144–153.

Carver, T., Harris, S. R., Berriman, M., Parkhill, J. & Mcquillan, J. A. 2011. Artemis: an integrated platform for visualization and analysis of high-throughput sequence-based experimental data. Bioinformatics, 28, 464–469.

Choudhury, P. & Flower, A. M. 2015. Efficient assembly of ribosomes is inhibited by deletion of bipA in Escherichia coli. J Bacteriol, 197, 1819–27.

Cotteret, C., Vallières, E., Roy, H., Ovetchkine, P., Longtin, J. & Bussières, J. F. 2016. Profil de consommation et de sensibilité aux antibiotiques utilisés dans un centre hospitalier universitaire : étude rétrospective sur cinq ans. Archives de Pédiatrie, 23, 1040–1049.

Drawz, S. M. & Bonomo, R. A. 2010. Three decades of beta-lactamase inhibitors. Clin Microbiol Rev, 23, 160–201.

Holden, E. R., Yasir, M., Turner, A. K., Wain, J., Charles, I. G. & Webber, M. A. 2023. Genome-wide analysis of genes involved in efflux function and regulation within Escherichia coli and Salmonella enterica serovar Typhimurium. Microbiology (Reading), 169.

Ikeuchi, Y., Shigi, N., Kato, J., Nishimura, A. & Suzuki, T. 2006. Mechanistic insights into sulfur relay by multiple sulfur mediators involved in thiouridine biosynthesis at tRNA wobble positions. Mol Cell, 21, 97–108.

Kammen, H. O., Marvel, C. C., Hardy, L. & Penhoet, E. E. 1988. Purification, structure, and properties of Escherichia coli tRNA pseudouridine synthase I. J Biol Chem, 263, 2255–63.

Kern, W. V., Oethinger, M., Jellen-Ritter, A. S. & Levy, S. B. 2000. Non-target gene mutations in the development of fluoroquinolone resistance in Escherichia coli. Antimicrob Agents Chemother, 44, 814–20.

Knothe, H., Shah, P., Krcmery, V., Antal, M. & Mitsuhashi, S. 1983. Transferable resistance to cefotaxime, cefoxitin, cefamandole and cefuroxime in clinical isolates of Klebsiella pneumoniae and Serratia marcescens. Infection, 11, 315–7.

Kong, K. F., Schneper, L. & Mathee, K. 2010. Beta-lactam antibiotics: from antibiosis to resistance and bacteriology. Apmis, 118, 1–36.

Naas, T., Oueslati, S., Bonnin, R. A., Dabos, M. L., Zavala, A., Dortet, L., Retailleau, P. & Iorga, B. I. 2017. Beta-lactamase database (BLDB) – structure and function. Journal of Enzyme Inhibition and Medicinal Chemistry, 32, 917–919.

O‘neill, J. 2014. Antimicrobial Resistance: Tackling a Crisis for the Health and Wealth of Nations: December 2014. https://amr-review.org/sites/default/files/AMR%20Review%20Paper%20-%20Tackling%20a%20crisis%20for%20the%20health%20and%20wealth%20of%20nations_1.pdf.

Oethinger, M., Kern, W. V., Jellen-Ritter, A. S., Mcmurry, L. M. & Levy, S. B. 2000. Ineffectiveness of Topoisomerase Mutations in Mediating Clinically Significant Fluoroquinolone Resistance in Escherichia coli in the Absence of the AcrAB Efflux Pump. Antimicrobial Agents and Chemotherapy, 44, 10–13.

Perry, C. M. & Markham, A. 1999. Piperacillin/Tazobactam. Drugs, 57, 805–843.

Piddock, L. J. 2006. Clinically relevant chromosomally encoded multidrug resistance efflux pumps in bacteria. Clin Microbiol Rev, 19, 382–402.

Schwartz, C. J., Giel, J. L., Patschkowski, T., Luther, C., Ruzicka, F. J., Beinert, H. & Kiley, P. J. 2001. IscR, an Fe-S cluster-containing transcription factor, represses expression of Escherichia coli genes encoding Fe-S cluster assembly proteins. Proc Natl Acad Sci U S A, 98, 14895–900.

Shimada, T., Yamazaki, Y., Tanaka, K. & Ishihama, A. 2014. The Whole Set of Constitutive Promoters Recognized by RNA Polymerase RpoD Holoenzyme of Escherichia coli. PLOS ONE, 9, e90447.

Song, Y., Zhou, Z., Gu, J., Yang, J., Deng, J. & Wang, H. 2021. Reducing the Periplasmic Glutathione Content Makes Escherichia coli Resistant to Trimethoprim and Other Antimicrobial Drugs. Microbiology Spectrum, 9, e00743–21.

Stevenson, G., Neal, B., Liu, D., Hobbs, M., Packer, N. H., Batley, M., Redmond, J. W., Lindquist, L. & Reeves, P. 1994. Structure of the O antigen of Escherichia coli K-12 and the sequence of its rfb gene cluster. J Bacteriol, 176, 4144–56.

Sukhodolets, M. V., Cabrera, J. E., Zhi, H. & Jin, D. J. 2001. RapA, a bacterial homolog of SWI2/SNF2, stimulates RNA polymerase recycling in transcription. Genes Dev, 15, 3330–41.

Suzuki, H., Nishimura, Y. & Hirota, Y. 1978. On the process of cellular division in Escherichia coli: a series of mutants of E. coli altered in the penicillin-binding proteins. Proc Natl Acad Sci U S A, 75, 664–8.

Thomson, N. M., Turner, A. K., Yasir, M., Bastkowski, S., Lott, M., Webber, M. A. & Charles, I. G. 2022. A whole-genome assay identifies four principal gene functions that confer tolerance of meropenem stress upon Escherichia coli. Frontiers in Antibiotics, 1.

Tokumoto, U. & Takahashi, Y. 2001. Genetic analysis of the isc operon in Escherichia coli involved in the biogenesis of cellular iron-sulfur proteins. J Biochem, 130, 63–71.

Tooke, C. L., Hinchliffe, P., Bragginton, E. C., Colenso, C. K., Hirvonen, V. H. A., Takebayashi, Y. & Spencer, J. 2019. β-Lactamases and β-Lactamase Inhibitors in the 21st Century. J Mol Biol, 431, 3472–3500.

Trampari, E., Holden, E. R., Wickham, G. J., Ravi, A., Martins, L. D. O., Savva, G. M. & Webber, M. A. 2021. Exposure of Salmonella biofilms to antibiotic concentrations rapidly selects resistance with collateral tradeoffs. npj Biofilms and Microbiomes, 7, 3.

Turner, A. K., Eckert, S. E., Turner, D. J., Yasir, M., Webber, M. A., Charles, I. G., Parkhill, J. & Wain, J. 2020a. A whole-genome screen identifies Salmonella enterica serovar Typhi genes involved in fluoroquinolone susceptibility. Journal of Antimicrobial Chemotherapy, 75, 2516–2525.

Turner, A. K., Yasir, M., Bastkowski, S., Telatin, A., Page, A., Webber, M. & Charles, I. 2021. Chemical biology-whole genome engineering datasets predict new antibacterial combinations. Microb Genom, 7.

Turner, A. K., Yasir, M., Bastkowski, S., Telatin, A., Page, A. J., Charles, I. G. & Webber, M. A. 2020b. A genome-wide analysis of Escherichia coli responses to fosfomycin using TraDIS-Xpress reveals novel roles for phosphonate degradation and phosphate transport systems. Journal of Antimicrobial Chemotherapy, 75, 3144–3151.

Valéria Dos Santos, K., Diniz, C. G., De Castro Veloso, L., Monteiro De Andrade, H., Da Silva Giusta, M., Da Fonseca Pires, S., Santos, A. V., Morais Apolônio, A. C., Roque De Carvalho, M. A. & De Macêdo Farias, L. 2010. Proteomic analysis of Escherichia coli with experimentally induced resistance to piperacillin/tazobactam. Research in Microbiology, 161, 268–275.

Vinella, D., Gagny, B., Joseleau-Petit, D., D‘ari, R. & Cashel, M. 1996. Mecillinam resistance in Escherichia coli is conferred by loss of a second activity of the AroK protein. Journal of Bacteriology, 178, 3818–3828.

Yasir, M., Turner, A. K., Bastkowski, S., Baker, D., Page, A. J., Telatin, A., Phan, M. D., Monahan, L., Savva, G. M., Darling, A., Webber, M. A. & Charles, I. G. 2020. TraDIS-Xpress: a high-resolution whole-genome assay identifies novel mechanisms of triclosan action and resistance. Genome Res.

