## Supplementary information for "Tazobactam selects for multidrug resistance"

### Supplementary Data

#### **Supplementary table 1**: Genes identified by TraDIS-*Xpress* to have a role in susceptibility of *E. coli* to piperacillin (Pip) and tazobactam (Tazo), either separately or in combination (PipTazo). Log_2_-fold change (Log_2_FC) values shown are statistically significant (*q* < 0.05, *p*-value corrected for false discovery rate) and measure the difference in insertion mutants between antibiotic-treated and untreated (ctrl) conditions. These values could not be calculated for areas outside of genes, and therefore are not shown where insertions denote a change in expression affects fitness. In these cases, the effect of the gene’s expression on fitness is describes in the ‘observed change’ column.

| **Pathway** | **Gene** | **Pip vs ctrl** |  |  | **Tazo vs ctrl** | **PipTazo vs ctrl** |  |
| --- | --- | --- | --- | --- | --- | --- | --- |
|  |  | **LogFC**  **(¼ / ½ / 1 / 2xMIC)** | **Observed change** | **LogFC**  **(½ xMIC)** | **Observed change** | **LogFC**  **(¼ / ½ / 1 / 2xMIC)** | **PipTazo vs ctrl**  **Observed change** |
| **Efflux pumps & efflux regulators** | *acrA* |  | Increased expression beneficial at sub-MIC concentrations | -1.8 | Fewer insertions & Increased expression beneficial at sub-MIC concentrations |  | Increased expression beneficial at sub-MIC concentrations |
|  | *acrB* |  |  | -1.1 | Fewer insertions & Increased expression beneficial at sub-MIC concentrations |  | Increased expression beneficial |
|  | *acrR* | 2.5 / 1.2 / - / - | More insertions at sub-MIC concentrations | 11.8 | More insertions | 12.2 / 12.0 / 1.0 / 1.8 | More insertions |
|  | *acrE* |  |  |  | Increased expression beneficial |  | Increased expression beneficial at sub-MIC concentrations |
|  | *acrF* |  |  |  | Increased expression beneficial |  | Increased expression beneficial at sub-MIC concentrations |
|  | *tolC* |  |  | -0.5 | Fewer insertions | -1.3 / - / -1.4 / -2.2 | Fewer insertions |
|  | *mprA* |  |  | 3.5 | More insertions |  |  |
|  | *mdtE* |  |  |  | Increased expression beneficial |  |  |
|  | *mdtF* |  |  | -1.6 | Fewer insertions & Increased expression beneficial |  |  |
|  | *mdfA* |  |  |  | Increased expression beneficial |  |  |
|  | *marR* |  |  | 14.9 | More insertions | 12.0 / 15.7 / - / - | More insertions |
|  | *marA* |  |  | -1.0 | Fewer insertions & Increased expression beneficial |  | Increased expression beneficial at sub-MIC concentrations |
|  | *soxR* |  |  | 15.3 | More insertions | 11.6 / 11.8 / - / - | More insertions at sub-MIC concentrations |
|  | *soxS* |  |  |  | Increased expression beneficial |  | Increased expression beneficial at sub-MIC concentrations |
|  | *rob* |  |  |  | Increased expression beneficial |  | Increased expression beneficial at sub-MIC concentrations |
| **Transport across membranes & outer membrane porins** | *chbB* | -1.0 / -1.2 / - / -1.0 | Fewer insertions |  |  |  |  |
|  | *trkA* | -1.1 / -/ -/ - | Fewer insertions at sub-MIC concentrations |  |  |  |  |
|  | *pitA* |  |  | 5.0 | More insertions | 3.1 / - / 0.9 / 1.7 | More insertions |
|  | *ompC* |  |  | -2.3 | Fewer insertions | -1.8 / - / -4.9 / -5.0 | Fewer insertions |
|  | *nhaA* |  |  | -2.1 | Fewer insertions |  |  |
|  | *sapF* |  |  | -0.5 | Fewer insertions |  |  |
|  | *sapD* |  |  | -1.9 | Fewer insertions |  |  |
|  | *ptsP* |  |  | 2.5 | More insertions |  |  |
|  | *ptsN* |  |  |  | Increased expression beneficial |  |  |
|  | *tatA* |  |  |  |  | - / - / 1.1 / 1.5 | More insertions at above MIC concentrations |
|  | *tatB* |  |  |  |  | - / - / 1.0 / 1.4 | More insertions at above MIC concentrations |
|  | *tatC* |  |  |  |  | - / - / 1.2 / 1.6 | More insertions at above MIC concentrations |
|  | *tatD* |  |  |  |  | - / - / -0.6 / -0.6 | More insertions at above MIC concentrations |
| **Extracellular polysaccharides (LPS/ ECA)** | *wecA* | - / -0.7 / -0.8 / -0.7 | Fewer insertions |  |  | - / - / -0.8 / -1.7 | Fewer insertions at above MIC concentrations |
|  | *wzzE* | - / -0.8 / -1.0 / -0.7 | Fewer insertions |  |  | - / - / -1.5 / -1.6 | Fewer insertions at above MIC concentrations |
|  | *wecB* | - / -0.6 / -0.7 / -0.6 | Fewer insertions |  |  | - / - / -1.1 / -1.6 | Fewer insertions at above MIC concentrations |
|  | *wecC* | - / -0.9 / -1.1 / -0.6 | Fewer insertions |  |  | - / - / -1.1 / -1.7 | Fewer insertions at above MIC concentrations |
|  | *wecD* |  |  |  |  | - / - / -1.8 / -2.3 | Fewer insertions at above MIC concentrations |
|  | *wecE* | -0.5 / -0.8 / -0.7 / -0.5 | Fewer insertions |  |  | - / - / -2.1 / -2.8 | Fewer insertions at above MIC concentrations |
|  | *wzxE* | -0.9 / -1.1 / -1.0 / -0.9 | Fewer insertions |  |  | - / - / -1.9 / -2.3 | Fewer insertions at above MIC concentrations |
|  | *wecF* |  |  |  |  | - / - / -1.7 / -2.3 | Fewer insertions at above MIC concentrations |
|  | *wecG* | - / -0.8 / - / - | Fewer insertions |  |  |  |  |
|  | *wbbK* | -0.9 / -0.8 / - / -0.9 | Fewer insertions |  |  |  |  |
|  | *waaR* |  |  |  |  | - / - / -1.5 / -2.1 | Fewer insertions at above MIC concentrations |
|  | *waaS* | -0.7 / -0.4 / -0.1 / -0.3 | Fewer insertions |  |  |  |  |
|  | *lpxL* |  |  |  | Increased expression beneficial |  |  |
|  | *lptA* |  |  |  |  |  | Increased expression beneficial at above MIC concentrations |
|  | *lptB* |  |  |  |  |  | Increased expression beneficial at above MIC concentrations |
|  | *lptC* |  |  |  |  |  | Increased expression beneficial at above MIC concentrations |
|  | *lptD* |  |  |  |  |  | Increased expression beneficial at sub-MIC concentrations |
| **Cell envelope synthesis** | *mrcA* |  |  |  |  | - / - / -1.4 / -1.9 | Fewer insertions at above MIC concentrations |
|  | *mrcB* | -2.3 / -2.2 / -2.6 / -2.5 | Fewer insertions |  |  |  | Increased expression beneficial |
|  | *lpoA* |  |  |  |  | - / - / -1.3 / -1.6 | Fewer insertions at above MIC concentrations |
|  | *lpoB* | -4.0 / -3.6 / -4.7 / -5.2 | Fewer insertions |  |  | - / - / -4.8 / -4.9 | Fewer insertions at above MIC concentrations |
|  | *ldcA* | -3.5 / -3.8 / -3.8 / -4.2 | Fewer insertions |  |  | - / - / -4.8 / -4.2 | Fewer insertions at above MIC concentrations |
|  | *nlpI* | -2.0 / -1.8 / -1.9 / -2.1 | Fewer insertions |  |  | -2.1 / - / -3.4 / -3.8 | Fewer insertions |
|  | *slt* | -3.6 / -3.9 / -3.7 / -4.3 | Fewer insertions |  |  | -1.7 / - / -5.2 / -4.8 | Fewer insertions |
|  | *mepS* | - / -1.8 / - / -2.4 | Fewer insertions |  |  |  |  |
|  | *mlaA* |  |  | -1.4 | Fewer insertions |  |  |
|  | *mlaB* |  |  | -1.2 | Fewer insertions |  |  |
|  | *mlaC* |  |  | -1.9 | Fewer insertions |  |  |
|  | *mlaD* |  |  | -1.3 | Fewer insertions |  |  |
|  | *mlaE* |  |  | -1.1 | Fewer insertions |  |  |
|  | *ampG* |  |  |  |  | - / - / -1.8 / -1.7 | Fewer insertions at above MIC concentrations |
|  | *rodZ* |  |  |  |  | - / - / -3.0 / -3.9 | Fewer insertions at above MIC concentrations |
| **Translation** | *truA* | 1.3 / 1.3 / 1.5 / 1.5 | More insertions | 2.6 | More insertions | 3.8 / - / 1.7 / 2.0 | More insertions |
|  | *typA/ bipA* | 0.8 / 0.9 / 1.2 / 1.3 | More insertions | 3.4 | More insertions | 1.3 / - / 1.3 / 2.2 | More insertions at above MIC concentrations |
|  | *mnmE* | 0.9 / 0.8 / 1.0 / 1.5 | More insertions | 3.3 | More insertions | 4.3 / - / 2.0 / 2.9 | More insertions |
|  | *mnmG* | - / - / 1.0 / 1.4 | More insertions at above MIC concentrations | 2.9 | More insertions | 4.2 / - / 2.0 / 3.0 | More insertions |
|  | *tusA* |  |  |  |  |  | Reduced expression beneficial at sub-MIC concentrations |
|  | *tusB* |  |  | 4.2 | More insertions | 6.2 / - / - / - | More insertions at sub-MIC concentrations |
|  | *tusC* |  |  | 3.3 | More insertions | 5.6 / - / - / - | More insertions at sub-MIC concentrations |
|  | *tusD* |  |  | 2.7 | More insertions | 5.1 / - / - / - | More insertions at sub-MIC concentrations |
|  | *tufB* |  |  | 2.3 | More insertions | 4.7 / - / - / - | More insertions at sub-MIC concentrations |
|  | *glyT* |  |  |  | Increased expression beneficial |  | Increased expression beneficial at sub-MIC concentrations |
|  | *fusA* |  |  |  | Increased expression beneficial |  | Reduced expression beneficial at sub-MIC concentrations |
|  | *rplK* |  |  |  | Increased expression beneficial |  |  |
|  | *thrT* |  |  |  |  | 8.9 / 6.2 / 3.1 / 3.6 | More insertions |
|  | *usg* |  |  |  |  | 2.8 / - / 1.1 / 1.2 | More insertions |
|  | *deaD* |  |  |  |  | 2.3 / - / 0.7 / 0.7 | More insertions |
|  | *prfC* |  |  |  |  | 1.2 / - / 2.6 / 2.6 | More insertions |
|  | *prmB* |  |  |  |  | - / - / 2.5 / 2.9 | More insertions at above MIC concentrations |
| **Two component signalling systems** | *cpxA* |  |  | 1.9 | More insertions | 5.6 / - / 0.4 / 0.9 | More insertions |
|  | *cpxR* |  |  | 3.1 | More insertions |  | Increased expression beneficial at sub-MIC concentrations |
|  | *phoP* |  |  | -0.6 | Fewer insertions | - / - / -1.1 / - | Fewer insertions |
|  | *phoQ* |  |  | -2.4 | Fewer insertions | -1.1 / - / -1.5 / -2.4 | Fewer insertions |
|  | *rcsB* | - / -0.9 / -1.6 / -1.6 | Fewer insertions |  |  |  | Increased expression beneficial at sub-MIC concentrations |
|  | *arcB* |  |  | -0.8 | Fewer insertions |  |  |
|  | *envZ* |  |  |  |  | -1.1 / - / -2.2 / -2.4 | Fewer insertions |
|  | *ompR* |  |  |  |  | - / - / -3.4 / -2.4 | Fewer insertions at above MIC concentrations |
|  | *mzrA* |  |  | 3.9 | More insertions |  |  |
|  | *nlpE* |  |  |  |  |  | Increased expression beneficial |
|  | *safA* |  |  |  |  |  | Increased expression beneficial at sub-MIC concentrations |
| **DNA housekeeping, replication & repair** | *rssA* | - / -1.1 / -0.9 / -1.0 | Fewer insertions |  |  |  |  |
|  | *dcd* | 2.3 / 2.4 / - / - | More insertions at sub-MIC concentrations |  |  |  |  |
|  | *maoP* |  |  | 3.2 | More insertions |  |  |
|  | *fis* |  |  |  | Increased expression beneficial |  |  |
|  | *ihfB* |  |  |  |  | - / - / -6.9 / -5.5 | Fewer insertions at above MIC concentrations |
|  | *xseB* |  |  |  |  |  | Fewer insertions |
|  | *mutT* |  |  |  |  |  | Increased expression beneficial |
|  | *uspB* |  |  |  |  |  | Increased expression beneficial |
| **Cell division** | *tolQ* |  |  |  |  | - / - / -5.1 / -3.2 | Fewer insertions at above MIC concentrations |
|  | *tolR* | -2.5 / -2.7 / -3.0 / - | Fewer insertions |  |  | - / - / -3.3 / -5.5 | Fewer insertions at above MIC concentrations |
|  | *tolA* | -3.5 / -3.3 / -3.2 / -3.5 | Fewer insertions |  |  | - / - / -5.0 / -4.7 | Fewer insertions at above MIC concentrations |
|  | *tolB* | -3.3 / -3.3 / -2.9 / -3.5 | Fewer insertions |  |  | - / - / -9.6 / -9.3 | Fewer insertions at above MIC concentrations |
|  | *pal* | -1.9 / - / - / - | Fewer insertions at sub-MIC concentrations |  |  | - / - / - / -7.5 | Fewer insertions at above MIC concentrations |
|  | *damX* |  |  | 1.9 | More insertions |  |  |
|  | *ftsE* |  |  |  |  | -1.8 / -3.0 / -2.1 / -2.1 | Fewer insertions |
|  | *ftsN* |  |  |  |  | - / - / -1.7 / -2.0 | Fewer insertions at above MIC concentrations |
|  | *ftsX* |  |  |  |  | - / - / -2.0 / -2.3 | Fewer insertions at above MIC concentrations |
|  | *minC* |  |  |  |  | -1.1 / - / -2.8 / -2.4 | Fewer insertions |
|  | *minD* |  |  |  |  | -1.2 / - / -2.3 / -2.0 | Fewer insertions |
|  | *minE* |  |  |  |  | - / - / -2.8 / -3.3 | Fewer insertions |
|  | *envC* |  |  |  |  | - / - / -1.2 / -0.9 | Fewer insertions at above MIC concentrations |
|  | *amiB* |  |  |  |  | 1.7 / - / - / - | More insertions at sub-MIC concentrations |
|  | *hflD* |  |  |  |  |  | Increased expression beneficial at sub-MIC concentrations |
| **Proteases** | *lon* | - / -0.9 / - / - | Fewer insertions at sub-MIC concentrations | 7.4 | More insertions | 4.7 / - / - / - | More insertions at sub-MIC concentrations |
|  | *prc* | -2.0 / -2.3 / -2.4 / -2.5 | Fewer insertions |  |  | - / - / -4.0 / -4.2 | Fewer insertions at above MIC concentrations |
|  | *smpB* | 2.1 / 2.1 / 2.6 / 2.3 | More insertions |  |  | - / - / 2.8 / 3.7 | More insertions at above MIC concentrations |
|  | *rhlB* |  |  | 2.7 | More insertions |  |  |
|  | *degP* |  |  |  |  |  | Increased expression beneficial |
| **Chaperones** | *dsbA* | - / - / 1.3 / 1.6 | More insertions at above MIC concentrations | 3.0 | More insertions | 8.6 / 5.3 / 2.3 / 2.8 | More insertions |
|  | *dsbB* |  |  |  |  | 3.0 / - / - / - | More insertions at sub-MIC concentrations |
|  | *surA* |  |  | -0.2 | Fewer insertions & Reduced expression beneficial |  |  |
|  | *ibpB* |  |  | -1.7 | Fewer insertions |  |  |
|  | *ibpA* |  |  | -0.6 | Fewer insertions |  |  |
|  | *tig* |  |  | -2.0 | Fewer insertions |  |  |
|  | *bepA* |  |  |  | Increased expression beneficial |  |  |
| **Transcription** | *rapA* | 1.3 / 1.2 / 1.1 / 0.9 | More insertions | 4.1 | More insertions | 8.6 / 4.2 / 1.1 / - | More insertions at sub-MIC concentrations |
|  | *greA* |  |  | 6.0 | More insertions |  |  |
|  | *nusA* |  |  |  | Increased expression beneficial |  |  |
| **Respiration, Electron transport & ATP synthesis** | *atpB* | - / - / -2.1 / - | Fewer insertions at sub-MIC concentrations &  Increased expression beneficial at above MIC concentrations |  |  |  | Increased expression beneficial |
|  | *pgi* |  |  | -1.2 | Fewer insertions |  |  |
|  | *cra* |  |  |  |  | 2.8 / - / 1.4 / 1.8 | More insertions |
|  | *cydB* |  |  |  |  | - / - / 2.2 / 2.1 | More insertions at above MIC concentrations |
| **Amino acid metabolism** | *aroK* |  |  | 10.6 | More insertions | 6.1 / 3.3 / - / 2.5 | More insertions |
|  | *dapF* | -1.7 / - / - / - | Fewer insertions at sub-MIC concentrations |  |  |  |  |
|  | *glnD* |  |  |  |  | 2.9 / - / 0.9 / 1.1 | More insertions |
| **Stress response transcription factors & regulation** | *rpoS* |  |  | -2.0 | Fewer insertions | -2.2 / - / -1.8 / -3.3 | Fewer insertions & Increased expression beneficial |
|  | *yheO* |  |  |  |  | 3.1 / - / - / - | More insertions at sub-MIC concentrations |
|  | *lrp* |  |  | 2.5 | More insertions |  |  |
| **Fe-S cluster synthesis & repair** | *rseC* |  |  | 3.2 | More insertions | 1.1 / - / - / - | More insertions at sub-MIC concentrations |
|  | *iscS* |  |  |  |  |  | Increased expression beneficial at sub-MIC concentrations |
|  | *iscR* |  |  |  |  | 4.4 / - / -1.2 / - | More insertions at sub-MIC concentrations |
|  | *hscA* |  |  |  |  | - / - / 2.0 / 2.4 | More insertions at above MIC concentrations |
|  | *hscB* |  |  |  |  | - / - / 2.9 / 3.0 | More insertions at above MIC concentrations |
| **Glutathione biosynthesis** | *gshA* |  |  | 8.4 | More insertions |  |  |
|  | *gshB* |  |  | 4.8 | More insertions |  |  |
| **Osmoregulated periplasmic glucans** | *opgG* |  |  | 5.4 | More insertions | - / - / 1.4 / 1.9 | More insertions at above MIC concentrations |
|  | *opgH* | -0.7 / - / - / - | Fewer insertions at sub-MIC concentrations | 4.9 | More insertions | - / - / 0.8 / 1.3 | More insertions at above MIC concentrations |
| **Toxin-Antitoxin systems** | *higA* | -2.5 / -2.5 / - / - | Fewer insertions at sub-MIC concentrations |  |  |  |  |
|  | *dinQ* | - / - / - / 2.2 | More insertions at above MIC concentrations |  |  |  |  |
| **Acid stress** | *yodD* |  |  | -3.6 | Fewer insertions |  |  |
|  | *ydeP* |  |  |  |  |  | Increased expression beneficial at sub-MIC concentrations |
| **Small regulatory RNAs** | *hfq* |  |  |  | Increased expression beneficial |  |  |
| **Motility** | *hdfR* |  |  | 2.9 | More insertions |  |  |
| **cAMP** | *cyaA* |  |  |  |  | 2.8 / - / - / - | More insertions at sub-MIC concentrations |
| **Fimbriae-like protein** | *ydeQ* |  |  |  |  |  | Increased expression beneficial at sub-MIC concentrations |
| **Unknown** | *ychJ* | - / -1.2 / -1.3 / -1.3 | Fewer insertions |  |  |  |  |
|  | *ybcK* |  |  |  | Increased expression beneficial |  |  |
|  | *ybcM* |  |  |  | Increased expression beneficial |  |  |
|  | *yidQ* |  |  | 3.5 | More insertions |  |  |
|  | *ycbC* |  |  |  |  | - / - / -2.2 / -2.6 | Fewer insertions at above MIC concentrations |
|  | *yobH* |  |  |  |  |  | Increased expression beneficial at sub-MIC concentrations |

**Supplementary figure 1:**

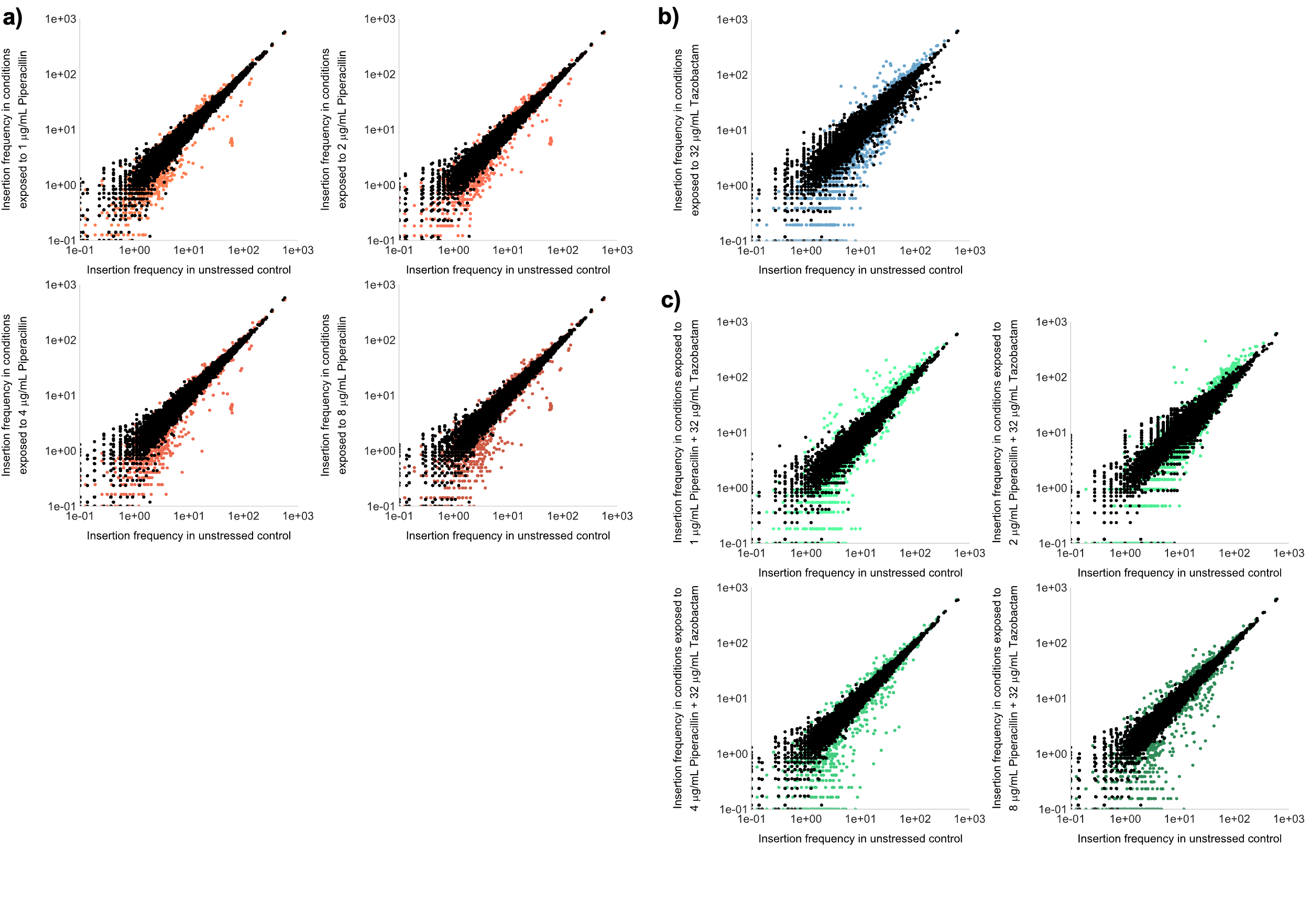

#### **Figure S1**: Insertion frequencies per gene for *E. coli* treated with **a)** piperacillin, **b)** tazobactam, and **c)** a combination of both drugs, relative to unstressed controls. Black points represent the insertion frequency per gene for each replicate to show variation between the replicates, and coloured points show the insertion frequency per gene between the control (x-axes) and each condition (y-axes).

**Supplementary table 2:** Single nucleotide polymorphisms (SNPs) identified in *Escherichia coli* and *Klebsiella* *pneumoniae* continuously cultured under increasing concentrations of piperacillin (Pip), tazobactam (Tazo) and a combination of the two (PipTazo). Genes in **bold** were also found in the TraDIS-*Xpress* analysis. Each condition consisted of three independent replicates, of which three single colonies were isolated and whole genome nucleotide sequence determined.

| **Species** | **Gene** | **SNP** | **Conditions** |
| --- | --- | --- | --- |
| *Escherichia coli* | *rpoD* | missense variant Asp445Ala | Pip & PipTazo |
|  | ***marR*** | frameshift variant Val63 | Pip & PipTazo |
|  |  | stop gained Trp83* | PipTazo |
|  |  | missense variant Val84Gly | PipTazo & Tazo |
|  | ***acrR*** | frameshift variant Arg168 | PipTazo |
|  | ***aroK*** | frameshift variant Arg120 | PipTazo |
| *Klebsiella pneumoniae* | ***cpxA*** | Missense Arg44Pro | Tazo only |
|  |  | Missense Leu30Gln | PipTazo only |
|  | ***acrR*** | conservative inframe deletion Leu179-Ala183 | Tazo only |
|  |  | disruptive inframe deletion Met109-Arg121 | PipTazo & Tazo |
|  | *rfbB* | frameshift variant Val220 | PipTazo only |
|  | *rfbD* | stop gained Tyr187* | Tazo only |
|  | *lldR* | missense variant Val60Gly | PipTazo only |
|  | *ftsI* | missense variant Arg167Ser | PipTazo & Tazo |
|  | ***ompC*** | conservative inframe deletion Ala365-Gly367 | Tazo only |
|  | *gltA* | missense variant Glu217Asp | Pip only |
|  | *cpdB* | frameshift variant Gly216 | Pip only |
|  | *mrdA* | frameshift variant Ala599 | Pip, Tazo & PipTazo |
|  | *tli1* | missense variant Pro13Leu | Pip, Tazo & PipTazo |
|  | Unknown gene  (homologous to BN427_0879 in ST258-K28BO) | frameshift variant Leu15 | Pip, Tazo & PipTazo |
